# PhyGCN: Pre-trained Hypergraph Convolutional Neural Networks with Self-supervised Learning

**DOI:** 10.1101/2023.10.01.560404

**Authors:** Yihe Deng, Ruochi Zhang, Pan Xu, Jian Ma, Quanquan Gu

## Abstract

Hypergraphs are powerful tools for modeling complex interactions across various domains, including biomedicine. However, learning meaningful node representations from hypergraphs remains a challenge. Existing supervised methods often lack generalizability, thereby limiting their real-world applications. We propose a new method, Pre-trained Hypergraph Convolutional Neural Networks with Self-supervised Learning (PhyGCN), which leverages hypergraph structure for self-supervision to enhance node representations. PhyGCN introduces a unique training strategy that integrates variable hyperedge sizes with self-supervised learning, enabling improved generalization to unseen data. Applications on multi-way chromatin interactions and polypharmacy side-effects demonstrate the effectiveness of PhyGCN. As a generic framework for high-order interaction datasets with abundant unlabeled data, PhyGCN holds strong potential for enhancing hypergraph node representations across various domains.

## Introduction

Hypergraphs are essential data structures adept at modeling complex multi-entity relationships. Conventional graph neural networks (GNNs), limited to pairwise interactions, cannot effectively capture the higher-order interactions inherent to hypergraphs. This shortfall led to previous methods [1–3] expanding hypergraphs to graphs using clique expansion [4]. More recent works [5–10] have adopted GNN structure such as graph convolutional networks (GCNs) and extended them to the hypergraph setting. HyperGCN [5] and HNHN [6], in particular, achieved state-of-the-art performances on hypergraph benchmark datasets. However, these supervised methods heavily rely on labeled nodes, which are often scarce in practical scenarios. In situations with insufficient labels, these methods fail to fully utilize the rich structural information of hypergraphs, leading to less effective and poorly generalizable node representations.

Self-supervised learning (SSL), an approach that extract meaningful knowledge from abundant unlabeled data to enhance model generalization [11, 12], presents a promising strategy to address these challenges. SSL creates pretext tasks from unlabeled data to predict unobserved input based on the observed part, enhancing the generalizability of models trained for computer vision (e.g., ImageNet pretraining [13, 14]) and natural languages (e.g., BERT [15]). While SSL methods have been proposed for GNNs [16–23], their application to hypergraph learning is underexplored. Noteworthy works [24–26] have targeted specific applications such as recommender systems, while others [27] propose pre-training GNNs on hypergraphs, primarily focusing on hyperedge prediction. A detailed discussion of related works can be found in **Supplementary Note** A.

Here, we introduce PhyGCN to address the urgent need for a general method that can effectively leverage unlabeled data from hypergraphs to enhance node representation learning. This self-supervised method extracts knowledge from the hypergraph structure using self-supervised tasks, generating robust node representations for diverse downstream tasks. Constructing self-supervised tasks in hypergraphs poses a unique challenge due to the variable sizes of hyperedges. We tackle this challenge by designing a self-supervised task that predicts masked hyperedges from observed ones and by incorporating an attention mechanism into our model architecture [28] to predict variable-sized hyperedges.

Link prediction, an effective pre-training task for traditional graphs, typically involves designing a neural network with fixed-length input to predict edges. Hyperedge prediction, however, is more challenging due to variable sizes. Simple average pooling of node embeddings has been found insufficient to model a hyperedge [29]. Although some works [24–27] have explored the potential of self-supervised learning on hypergraphs, our PhyGCN method is the first straightforward and effective approach to utilize hypergraph structure and can be generally applied to many tasks.

We demonstrate the effectiveness of PhyGCN through various evaluations across multiple tasks and datasets, showing its advantage over state-of-the-art hypergraph learning methods. Notably, PhyGCN has been applied to study multi-way chromatin interactions and polypharmacy side-effect network data, confirming its advantages in producing enhanced node embeddings and modeling higher-order interactions. Together, PhyGCN, a self-supervised method for hypergraph representation learning, can be applied broadly to a variety of problems.

## Results

### Overview of PhyGCN

**Fig. 1** provides a detailed depiction of PhyGCN’s model architecture and general workflow. PhyGCN aims to enhance node representation learning in hypergraphs by effectively leveraging abundant unlabeled data. To this end, we propose 1) a self-supervised learning strategy that pre-trains the model directly on the hypergraph structure before proceeding with any downstream tasks, and 2) a corresponding model architecture comprising a base hypergraph convolutional network for representation learning, and an attention network [28] for executing the self-supervised task.

**Figure 1:**
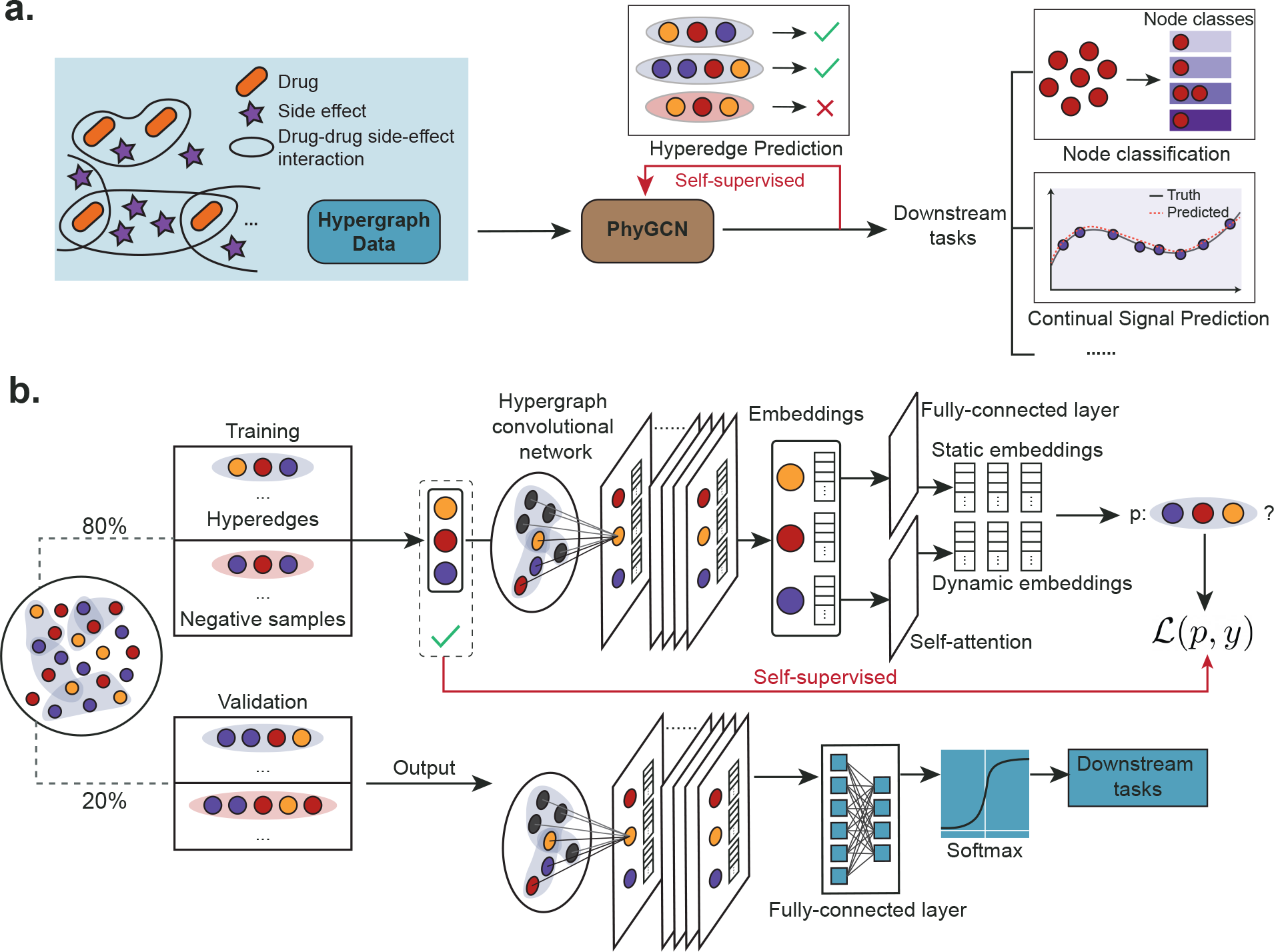
Overview of PhyGCN. **a**. The general workflow of PhyGCN begins with a hypergraph and its hyperedge information. Hyperedge prediction is employed as a self-supervised task for model pre-training. The pre-trained embedding model is then used for downstream tasks. **b**. This illustration elaborates on PhyGCN’s detailed workflow, which involves generating labeled data for self-supervised learning by randomly taking 80% of the hyperedges as positive samples, and creating negative samples by changing the nodes in each positive sample. This data facilitates the training of the base hypergraph convolutional network with the attention network. The pre-trained base model is subsequently used for downstream tasks with a fully connected layer.

As shown in **Fig. 1a**, to pre-train the model, we establish a self-supervised hyperedge prediction task using randomly masked hyperedges (20%). The neural networks are then trained to reconstruct them using only the remaining hyperedges (80%). The pre-trained neural network can then be used for downstream tasks such as node classification and continual signal prediction. See **Methods** for the details of the pre-training scheme.

**Fig. 1b** details the architecture of the neural network model, consistent of a base hypergraph convolutional network and two end networks for different tasks. The base convolutional network computes the embedding of each node in the hypergraph, which is then directed either to the attention network for variable-sized hyperedge prediction, or to a fully connected layer for node-level downstream tasks. During the pre-training phase, the base model and the attention network are trained while the fully connected layer for downstream tasks remains randomly initialized. We select and retain the base model that exhibits the best performance on the pre-training task (hyperedge prediction) on the validation set, and subsequently fine-tune it with the fully connected layer for downstream tasks. During this stage, we incorporate the original hypergraph data with its complete hyperedge information, and use the downstream task labels as the supervisory signals for training.

To calculate the embedding for a target node, the hypergraph convolutional network aggregates information from neighboring nodes connected to it via hyperedges, and combines it with the target node embedding to output a final embedding. These computations are carried out at each layer of the network. The base hypergraph convolutional network is a stack of such hypergraph convolutional layers (see **Methods**). To mitigate over-smoothing [30] and enhance the performance of the base model, we concatenate the output of each layer in the end to generate the final embeddings for the nodes. Furthermore, we adapt DropEdge [31] and introduce DropHyperedge, where we randomly mask the values of the adjacency matrix for the base hypergraph convolutional network during each training iteration to prevent overfitting and improve generalization in the pre-training phase. By integrating the model architecture and pre-training scheme, PhyGCN effectively learns from a hypergraph structure and applies the learned knowledge to downstream tasks.

### Self-supervised learning improves performance

We start with the fundamental question: *can PhyGCN more effectively utilize the structural information in a hypergraph?* To assess this, we benchmarked PhyGCN against the current state-of-the-art hypergraph learning methods, including HyperGCN [5] and HNHN [6]. We also established a baseline by merely decomposing the hypergraphs into graphs and applying a standard GCN [32]. We evaluated these methods using the benchmark node classification task on citation networks (including Citeseer, Cora, DBLP, and PubMed) whose hyperedges denote either co-authorship or co-citation relationships. Given that the baselines utilize different data splits for their evaluations, we consistently evaluated our method along with all the baselines on the same random data splits across a range of training data ratios. Detailed statistics, experiment settings, and results are provided in **Supplementary Note** B.2.

In order to ascertain how each hypergraph learning method performs with limited training data, we first set the training data ratio to 0.5% across the five datasets. The results are shown in **Fig. 2a**. Across 10 random data splits, we found that both HyperGCN and HNHN fail to outperform the standard GCN model, which merely transforms the hypergraphs into conventional graphs. Conversely, PhyGCN, equipped with pre-training on the self-supervisory signals obtained directly from the hypergraph, performs significantly better when dealing with a small training data ratio. It outperforms GCN across all datasets, particularly on small networks such as Citeseer and Cora. The improvement over GCN suggests that PhyGCN in more adept at learning from the higher-order hypergraph structure as opposed to merely decomposing it into a binary graph. Furthermore, the pre-training scheme allows PhyGCN to extract more information for node representation learning, especially when dealing with small hypergraphs.

**Figure 2:**
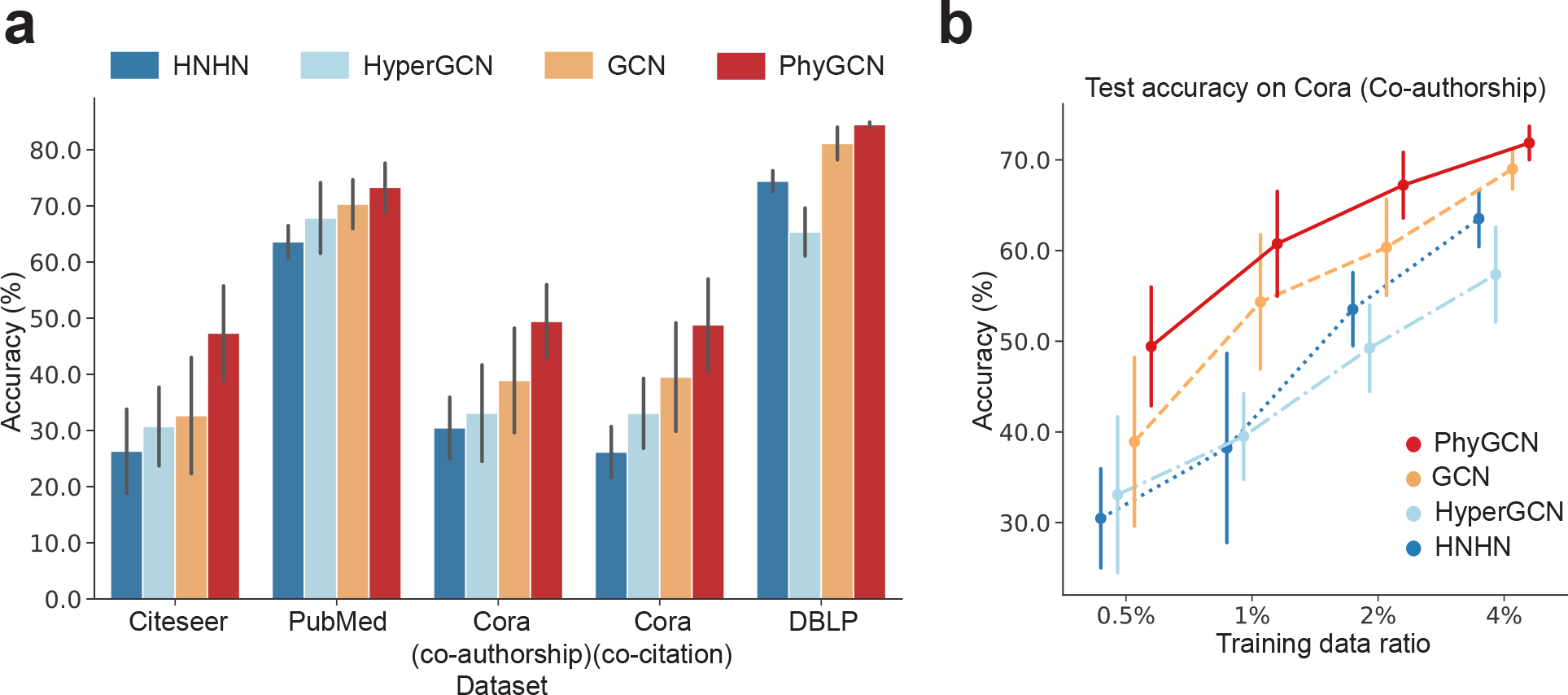
Evaluation of our model on node classification on citation networks with variable hyperedge sizes (Citeseer, Cora (co-authorship), Cora (co-citation), DBLP, and PubMed). The baseline methods include HNHN [6], HyperGCN [5], and vanilla GCN [32]. **a**. Comparison of PhyGCN’s accuracy with the baseline methods when the training data is minimal (0.05%). **b**. On the Cora (co-citation) dataset, PhyGCN consistently outperforms the baselines across various training data ratios (0.05%, 1%, 2%, 4%).

To evaluate PhyGCN against the baselines on a wider range of training data ratios, we generated data splits with increasing training data ratios of 1%, 2%, and 4%. The results on the Cora (co-authorship) dataset are shown in **Fig. 2b**, while comprehensive results of all models on the five datasets are provided in **Supplementary Note** B.2. Overall, PhyGCN outperforms the other methods, demonstrating more robust performance under random data splits. Across varying training data ratios, PhyGCN consistently outperforms the three baselines. At training data ratios of 2% and 4%, PhyGCN’s performance is considerably more stable than that of the baselines, as indicated by the significant decrease in the variance of PhyGCN’s accuracy across 10 random data splits.

These observations suggest that the pre-training stage enables the model to more effectively gather information from unlabeled data, allowing it to perform well even when a small amount of data is available. The results underscore the potential of PhyGCN in learning with higher-order interactions without the strong requirement of ample labeled data for a specific task. By fully leveraging the hypergraph structure for self-supervised training and designing a simple but effective model architecture, PhyGCN can achieve superior and more stable performances on downstream tasks compared to existing baselines.

### Exploring multi-way chromatin interaction

To further demonstrate the advantages of PhyGCN, particularly its potential for advancing discoveries in biological data, we applied it to multi-way chromatin interaction datasets. Such datasets reflect simultaneous interactions involving multiple genomic loci within the same nuclei, providing an opportunity to probe higher-order genome organization within a single nucleus. However, due to the limitation of current mapping technology and the exponentially growing combination space of genomic loci, these datasets tend to be extremely sparse and noisy. We evaluated our method on a node classification problem on the SPRITE [33] data from the GM12878 cell line, where the aim is to predict the Hi-C subcompartment label for each genomic bin. The details of the experimental setup, including the tasks, baselines, and datasets, can be found in **Supplementary Note** B.3.

#### Why pre-training?

We first evaluated how the pre-training scheme improves the learned representations for the downstream tasks with a node classification problem on the SPRITE data. In this task, we aim to predict the Hi-C subcompartment label for each genomic bin and we split the data by chromosome indices. **Fig. 3a** shows the results of the node classification problem. Without pre-training, our “plain” model is directly trained on the subcompartment labels to make predictions. With self-supervised learning that extracts the hyperedge information, the pre-trained model performs much better on the cross-chromosome subcompartment label prediction task.

**Figure 3:**
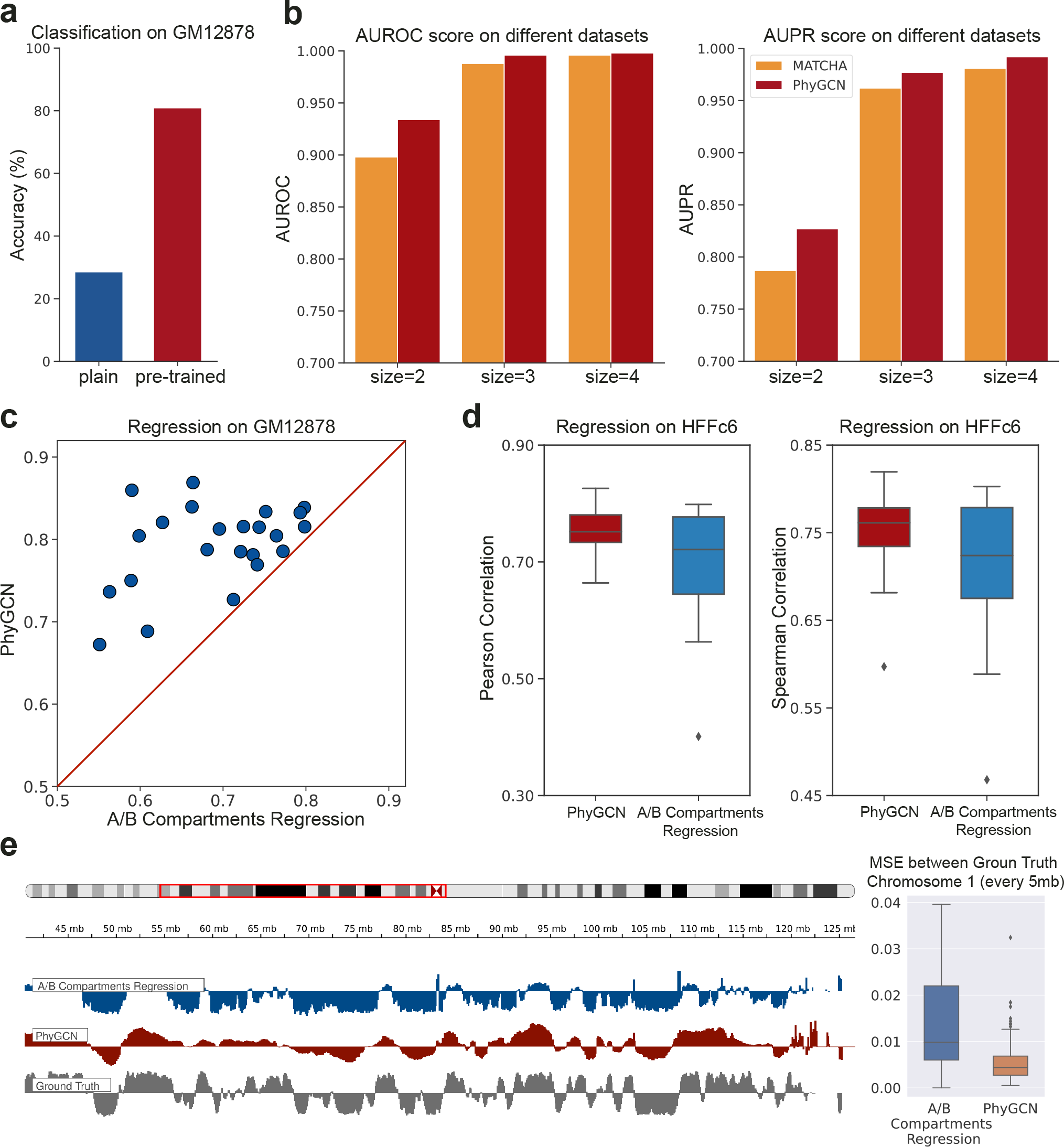
Application to the SPRITE data of chromatin interactions. **a**. Classification accuracy for subcompartment labels of the GM12878 cell line. **b**. AUROC and AUPR scores for predicting multi-way chromatin interactions, where we compare our method with MATCHA [34] across hyperedges of sizes 2, 3, and 4. **c**. Regression results on the GM12878 cell line, compared against the baseline A/B Compartments. **d**. Regression results on the HFFc6 cell line, where we compared our transferred PhyGCN results with the baseline A/B Compartments. **e**. Browser shot on chromosome 1 for Repli-seq prediction task (regression on HFFc6). The boxplot shows the MSE distribution between the ground truth and the result from PhyGCN or the baseline. MSE is calculated within each region of 5 Mb.

#### Why the hypergraph convolutional network?

In **Fig. 3b**, we investigate the effectiveness of PhyGCN’s hypergraph convolutional architecture in learning multi-way chromatin interactions of variable sizes, which is the pre-training task of the model. Specifically, we compared PhyGCN against MATCHA [34], which uses an autoencoder to generate node representations. As in the same setting in [34], we consider hyperedges with the size of 2, 3, and 4 and use the AUROC and AUPR scores as the evaluation metrics. We found that, with the hypergraph convolutional architecture, PhyGCN outperforms MATCHA on the three different hyperedge sizes with a small fraction of training data provided. It should be noted that the hyperedge prediction task aims to provide a pre-trained model for downstream tasks. To further evaluate the effectiveness of our base model, we performed several other hyperedge prediction tasks as a thorough ablation study. The results, detailed in **Supplementary Note** B.5, confirm that the introduced convolutional architecture enables the model to consistently and stably capture hyperedge information across different datasets, providing a better pre-training stage.

#### Comparison with previous methods

To investigate whether our pre-trained model captures informative patterns of multi-way chromatin interactions, we conducted a downstream regression task to predict the DNA replication timing of each genomic bin using corresponding embeddings as input. We fit the signals on even or odd chromosomes given the training labels on odd or even chromosomes. As a baseline, we used A/B compartment scores [35]. The results are shown in **Fig. 3c**, where the *x*-axis represents the Pearson correlation score of A/B compartment scores and Repli-seq signals, and the *y*-axis denotes the Pearson correlation score of the output of PhyGCN and ground truth labels. Each data point on **Fig. 3c** corresponds to a chromosome. Based on the downstream tasks, we found that with pretraining on the hyperedge interactions, our model can effectively capture the underlying patterns of 3D genome organization, generating more informative embeddings that are relevant to important biological functions such as DNA replication.

#### Transferring knowledge to a different hypergraph structure

We further explored the potential of PhyGCN on learning and adapting to new hypergraph structures. Specifically, we evaluated our method on transferring the association between multi-way chromatin interactions and DNA replication timing learned from the SPRITE data of the GM12878 cell line to that of the HFFc6 cell line. To achieve this, we first pre-trained our base model and the attention network to predict multi-way chromatin interactions of the GM12878 cell line using the self-supervised learning task. We then fixed the base model and fine-tune the fully connected layer to predict replication timing of genomic loci in the GM12878 cell line. Next, with the attention network fixed, we fine-tuned the base model with the task of predicting multi-way chromatin interactions in the HFFc6 cell line using the hypergraph constructed from the SPRITE data of HFFc6 following the same SSL setting as the dataset of the GM12878 cell line. With an additional regularizer on the base model weights, we expect our base model to learn the subtle knowledge in hypergraph structural differences and combined with the fixed fully connected layer to make accurate predictions of Repli-seq signals in HFFc6 cell line. The results are shown in **Fig. 3d**, where we use boxplots to show the Pearson and Spearman correlation scores on each chromosome and compare them with the A/B compartment score calculated from the contact of SPRITE data in HFFc6. In **Fig. 3e**, we show an example from chromosome 1 where we compared the ground truth Repli-seq signals to the predictions from A/B compartments regression and PhyGCN with PhyGCN reaching an overall smaller MSE. Together, our results demonstrate that PhyGCN can better learn the representation based on multi-way chromatin interactions and utilize it to make downstream inferences.

### Application to predicting polypharmacy side effect

Next, we sought to apply PhyGCN on the polypharmacy side effect dataset and demonstrate the advantages of learning such data as a hypergraph rather than a graph. Typically, the association between drugs and side effects is represented as a pairwise drug-side-effect network. However, during disease treatment, multiple drugs are often used in combination, and undesirable combinations can cause side effects not known to either individual drug in the combination (polypharmacy side-effect) [36]. Recent works like [36, 37] have used GNNs to predict polypharmacy side-effects with noteworthy performance. These methods treat drugs as nodes and side effects associated with a drug pair as the attributes for the corresponding edge. However, the number of unique side effects is comparable to or even exceeds the number of unique drugs, making it inefficient to model the data as multiple graphs for each side effect. Furthermore, real-world data likely involve drug combinations with more than two drugs that cause side-effects, which GNN methods may not be able to handle. In contrast, hypergraph methods like PhyGCN can easily adapt to such datasets. Since the polypharmacy side-effect data involves interactions beyond pairwise, it can be naturally learned as a hypergraph with drugs and side effects as the nodes. We investigated how PhyGCN learns such interactions and compared it with state-of-the-art methods, including graph-based methods such as Decagon [36] and ComplEX [37], and hypergraph-based method Hyper-SAGNN [28]. Further details are provided in **Supplementary Note** B.4.

In [36, 37], only side-effects known to be associated with more than 500 drug combinations were used for test data. In addition to this original setting, we report results on random data splits where any side effect can be used for testing. Our findings, presented in **Fig. 4a**, indicate that modeling the data as a hypergraph (using Hyper-SAGNN [28] and PhyGCN) yields better performance than modeling it as graphs (using Decagon [36] and ComplEX [37]) in both the original and random settings. We conducted additional experiments to investigate how PhyGCN benefits from learning the data as a hypergraph by testing it on different types of data splits. Specifically, we considered interactions that involve a known pairwise relationship as “expected” and those that do not involve any known relationship as “unexpected” in the original dataset, as some side-effects are already known to be associated with a single drug (see **Supplementary Note** B.4). We empirically studied how the baselines and our method performed when testing only on expected or unexpected data. As shown in **Fig. 4a**, GNN methods such as Decagon and ComplEX showed a drop in performance from “expected” to “unexpected”, while PhyGCN maintained a consistent performance on both types of data. As the unexpected polypharmacy side-effect triplet does not contain any potential pairwise interactions (while “expected” ones have), the “unexpected” test data serves as a useful indicator of how well a method captured such high-order interactions. The consistency of PhyGCN and Hyper-SAGNN on the different data splits further demonstrates the advantage of modeling complex interactions as a hypergraph. Additionally, our results showed that PhyGCN outperformed Hyper-SAGNN, indicating its superior capability to learn from hypergraphs.

**Figure 4:**
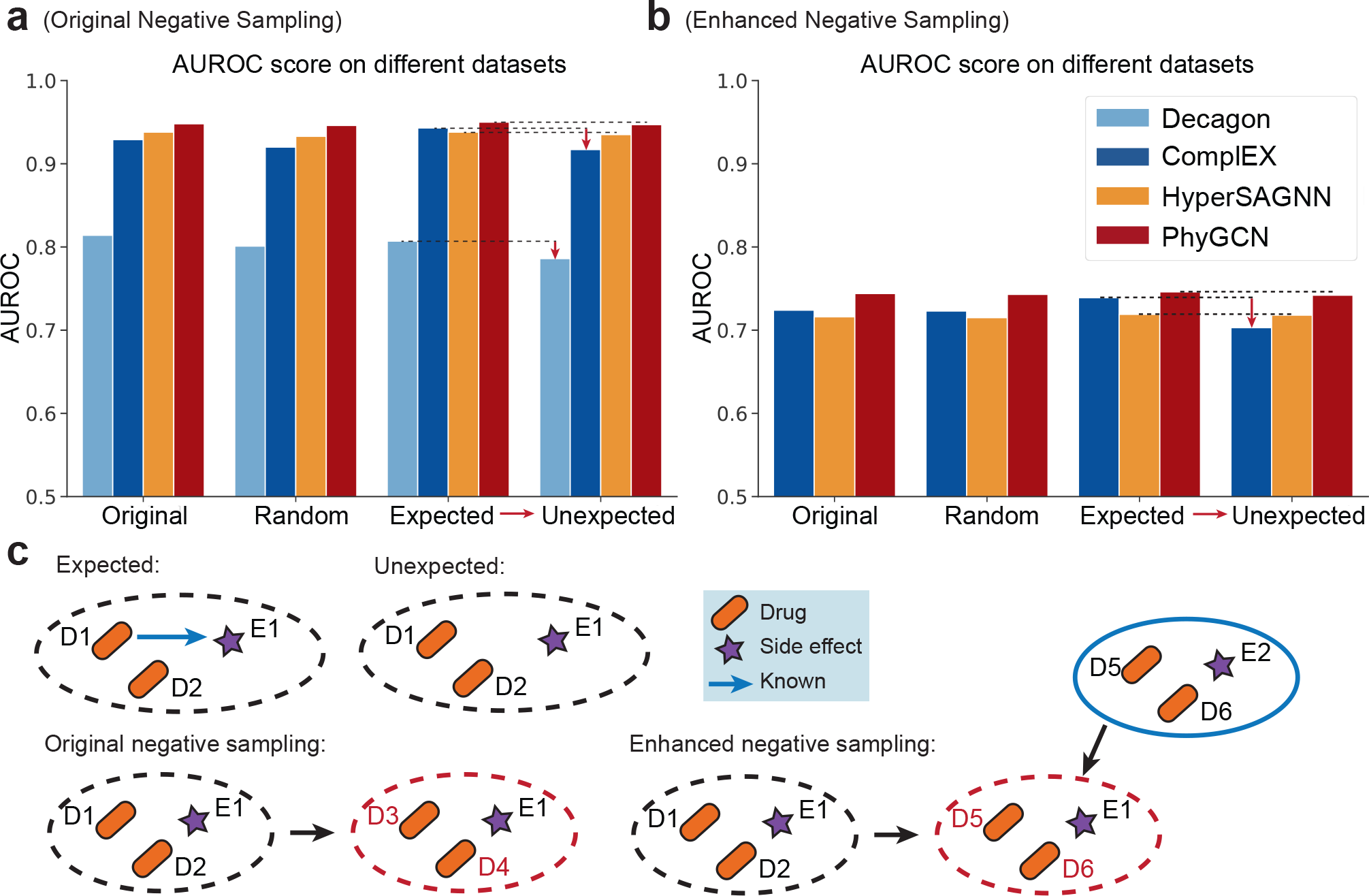
Application to the polypharmacy side effects dataset. **a**. We display the AUROC score for polypharmacy side effect prediction across different data splits, distinguishing between performances on “expected” and “unexpected” polypharmacy side effects. **b**. The AUROC score for polypharmacy side effect prediction across different data splits under an enhanced negative sampling strategy. **c**. We concretely demonstrate our definition of expected/unexpected relations. In the expected interactions, we have prior knowledge that one of the drugs causes the side effect, as indicated by the red arrow. Furthermore, we illustrate the difference between the original negative sampling and the enhanced negative sampling. The latter selects drug pairs that generate other side effects to be combined with the current example.

#### Enhanced negative sampling strategy

To investigate the effect of negative sampling on the polypharmacy side effect dataset, we conducted an experiment to compare the traditional random negative sampling method with our proposed negative sampling strategy. The traditional approach [36, 37] generates negative samples by randomly sampling two drugs that do not cause the specific side effect in the training data. However, we hypothesize that this strategy may lead to a model that learns the bias of the likelihood that a drug combination causes any side effect rather than the specific side effect in question. To address this, we propose a stronger criterion for negative sampling, which involves sampling a drug combination that is known to cause other side effects and combining it with the given side effect as a negative sample. This strategy forces the model to differentiate between the positive and negative samples and learn the subtle differences between them. In other words, the model is forced to make distinctions for each side effect. The results of our experiment are presented in **Fig. 4b**. We found that our proposed negative sampling strategy poses a greater challenge for all methods to perform well, with an average test accuracy below 80%. However, we observed that PhyGCN outperforms the baselines and performs stably well across different data splits. This strongly suggests that PhyGCN is better able to grasp the complex interactions without bias, unlike the baselines. We further present detailed examples in **Fig. 4c** to demonstrate the expected and unexpected side effect, and our enhanced negative sampling strategy. Together, the results of this experiment highlight the importance of careful negative sampling in achieving good performance in learning hypergraphs.

## Discussion

In this paper, we introduced PhyGCN, a hypergraph convolutional network model with a self-supervised pre-training scheme, designed to effectively capture higher-order information of hypergraphs. We demonstrated the effectiveness of PhyGCN on node classification tasks of benchmark citation networks and conduct comprehensive studies on a multi-way chromatin interaction dataset, highlighting the contribution of each component to capturing informative interaction patterns. We also showed the advantages of modeling polypharmacy side effects as hypergraphs and propose an improved negative sampling scheme to evaluate models with less bias.

While self-supervised training has been extensively studied for graphs with pairwise links [16–23], few works have explored it for hypergraph representation learning [27]. Our focus is on leveraging the attention mechanism [28] to predict hyperedges with arbitrary lengths and capture the indecomposiblity of a hyperedge, allowing PhyGCN to extract useful information from complex hypergraph structures.

With its ability to model higher-order interactions in various graph-structured data, such as social networks with group interactions, PhyGCN has great potential for a wide range of applications. Many real-world graph-structured data possess higher-order interactions. For example, social networks have group interactions that are inherently hypergraph-structured, and decomposing them into pairwise interactions may lose valuable information. By modeling such graph data as hypergraph, PhyGCN can provide valuable node representations by extracting information from the structure. This offers opportunities for leveraging the method in various applications, from drug discovery to social network analysis, where complex, multi-way interactions play a critical role.

## Methods

### Preliminaries

#### Graph Neural Networks

We denote a graph by *G* = (*V, E*), where *V* is the vertex set with node *v ∈ V* and *E* is the edge set with pairwise edge (*u, v*) *∈ E*. Graph neural networks (GNNs) mostly follow the following scheme [38]:

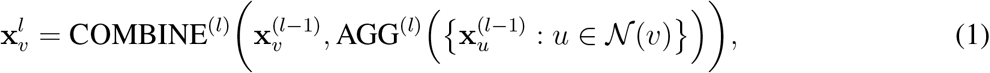

where 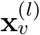 is the embedding vector of node *v* at layer *l* of the network, 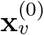 is the input data, and *𝒩*(*v*) is the set of nodes that are adjacent to *v*. Here AGG(*·*) and COMBINE(*·*) are two functions usually used to aggregate the neighboring embeddings and combine the aggregated embedding with the original node embedding to output the embedding of a given node. Different GNN models may have different choices of COMBINE(*·*) and AGG(*·*) functions. As a most popular approach to aggregate and combine the node vectors, GCN [32] averages over all the first-order neighboring nodes including the node itself. Let **X**^(*l*)^ represent the embedding matrix of all nodes at layer *l*, and the equation boils down to

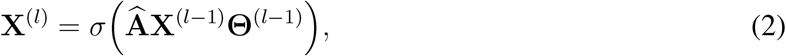

where *σ* is some activation function, 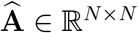 is the normalized adjacency matrix with N denoting the number of nodes. 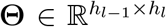 is the weight matrix at layer *l −* 1 where *h*_*l*_ as the hidden embedding size at each layer *l*. And 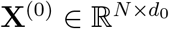 is the input node features of all nodes where *d*_0_ is the size of the input node feature size.

#### Hypergraph Convolutional Networks

In an attempt to adapt GCN for tasks on hypergraph, the problem naturally arises: how do we appropriately adapt the adjacency matrix for hypergraphs? Adjacency matrix is natural to construct for graphs: setting the entry **A**_*ij*_ = 1 for each edge (*v*_*i*_, *v*_*j*_). However, hyperedge involve more than two nodes and constructing **A** will thus be tricky. As [39] proposed, for a hypergraph *G* = (*V, E*), a hypergraph can be represented by an incidence matrix **H** *∈* R^*N ×M*^, with *N* as the number of nodes and *M* as the number of hyperedges. When a hyperedge *e*_*j*_ *∈ E* is incident with a vertex *v*_*i*_ *∈ 𝒱*, **H**_*ij*_ = 1, otherwise equals 0. We denote **W** as the hyperedge weight matrix, and then a straightforward definition for the adjacency matrix will be **A** = **HWH**^⊤^. And to normalize it similarly as in GCN, we take

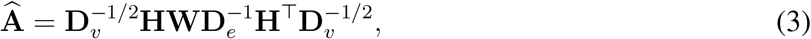

where the diagonal matrices **D**_*e*_ *∈* R^*M ×M*^, **D**_*v*_ *∈* R^*N ×N*^ respectively represent the degree of hyperedges and the degree of vertices [39] and 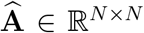. A layer of such hypergraph GCN model will then be similar to the regular GCN but with a different adjacency matrix

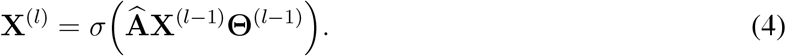

Similarly, 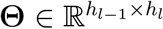 is the weight matrix at layer *l −* 1 and 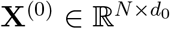 is the input node features of all nodes. We adopt such simple architectures in our work, along with several techniques in GNNs to enhance the model’s learning on both self-supervised and main tasks.

### Our Method

#### Base Hypergraph Convolutional Network

We thus propose a self-supervised learning strategy, along with a simple but effective hypergraph convolutional network architecture, to better capture the structural knowledge of a hypergraph. Our base convolutional model follows from the previous setup of a hypergraph convolutional network, where

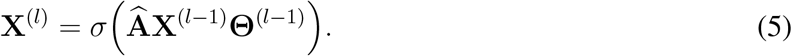

And we further introduce the following modifications to the base hypergraph convolutional model. **Skip/Dense Connection**. GCNs have been known to the have vanishing gradient problem where stacking multiple convolution layers in a GCN model causes the derivative of the loss function to approach zero and thus hard to train [40]. To make GCNs deeper and prevent the vanishing gradient problem, [41] proposes several adapted strategies based on the methods frequently used for CNNs. Similar vanishing gradient problems have also been observed in hypergraph convolutional networks [42]. Therefore, we design a similar but more computing-efficient strategy where we concatenate the outputs of all layers of the model only after the last layer (instead of concatenating the previous outputs at *each* layer as in [41]), as illustrated in Figure 5a. That is, we keep all the previous layers unchanged and set the last layer of the convolutional model as:

**Figure 5:**
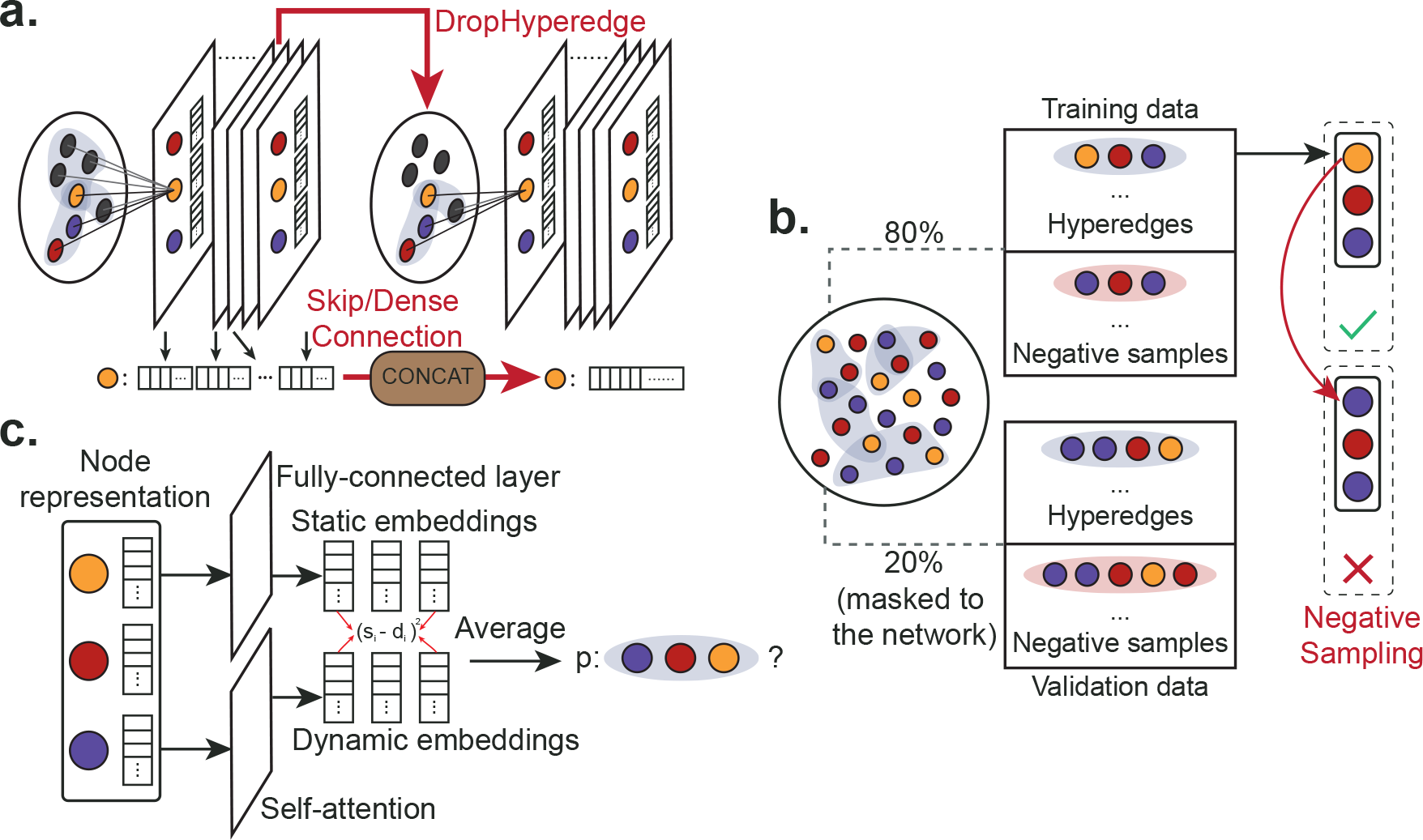
Illustration of the method. **a**. For the base hypergraph convolutional network, we propose two adapted strategies (DropHyperedge and Skip/Dense Connection) to further improve the pre-training of PhyGCN. **b**. For the self-supervised pre-training task, we randomly mask 20% of the hyperedges in the hypergraph and generate negative samples with regard to each positive training example. **c**. The pre-training scheme requires prediction of hyperedges that are arbitrarily sized. Therefore, we adopt an attention network [28] to fulfill the task.

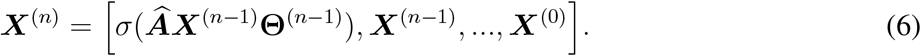

And we denote 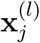 as the *j*-th row of **X**^(*l*)^, which is the embedding at layer *l* for node *j*. Such design helps us save computation time while preventing the model’s performance from dropping with more layers. The eventual base model we use is then the *n*-layer convolutional model with concatenation at the last layer. We either add an attention layer for the self-supervised task or an MLP layer for the main task.

### DropHyperedge

Overfitting is a common issue in the use of neural networks, where the network learns to fit the training data too well and fails to generalize to test data. To mitigate such an effect, dropout has been a popular technique for many fields and is also commonly used for GCNs. To further reduce overfitting and improve generalization in GCNs, [31] introduced a new regularization technique designed for GCNs called DropEdge, which randomly mask the edges of the graph at each iteration. In this work, to address the over-fitting issues with PhyGCN and encourage better generalization in the pre-training stage, we adapt the DropEdge strategy to our model by randomly masking the values on the adjacency matrix 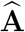 at each iteration (as illustrated in Figure 5a). Note that such a strategy is only used for the self-supervised hyperedge prediction task. We use the same dropout strategy in the main task as the other baselines.

#### Pre-training with Self-supervisory Signals

##### Self-supervised task

As shown in Figure 5b, given a hypergraph *G* = (*V, E*), we randomly divide the set of hyperedges *E* into the positive training set *E*_train_ and the positive validation set *E*_valid_. Moreover, we generate 5 negative samples for each hyperedge in 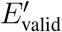 and 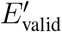. The final training and validation set is then

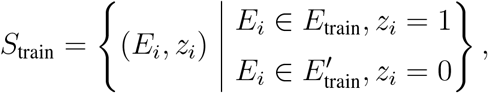

and

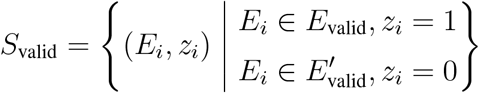

where *E*_*i*_ = *{v*_1_, …, *v*_*k*_*}* is a set of nodes of arbitrary size *k*. We then train the model by the hyperedge prediction task: given a set of node features 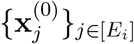 with arbitrary size *k* and 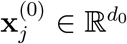, the goal is to predict *z*_*i*_ *∈ {*0, 1*}*: whether the nodes form a hyperedge. Therefore, given the training set *S*_train_, we aim to minimize the following cross-entropy loss with regard to each training batch *S*:

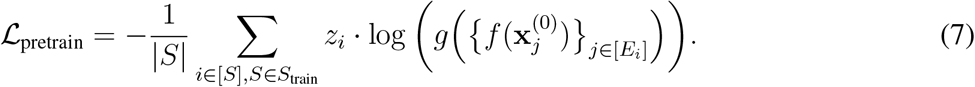

Here, 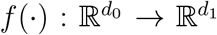 denotes the base hypergraph convolution model, where *d*_0_ is the input feature size and *d*_1_ is the size of the output embedding 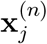 of the network. Moreover, 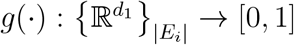 denotes the attention network, which takes a arbitrary-size set of node embeddings 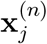 and output the probability *p* of whether the nodes form a hyperedge. The base hypergraph convolutional model *f* (*·*) and attention network *g*(*·*) are trained together, and the base model’s weight parameters are saved for further fine-tuning in downstream tasks.

##### Negative sampling

We generate negative samples similarly to [28, 29], where the negative samples are 5 times the number of positive samples provided by the hypergraph structure. Specifically, given a existing hyperedge *E*_*i*_ = *{v*_1_, …, *v*_*k*_*} ∈ E*_train_, we generate 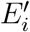 by randomly alternating one or multiple of the node in *E*_*i*_ such that the new set does not belong to the existing hypergedge set, as shown in Figure 5b. Moreover, the sampling strategy varies for different hypergraphs. For hypergraph with only one node type (for example, citation network), we have

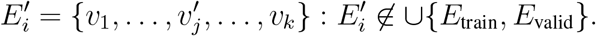

For hypergraphs with different node types (for example, polypharmacy side-effect data), we randomly alternate a node within its node type. Take hypergraph with two node types *v* and *u* for example, the hyperedge is *E*_*i*_ = *{v*_1_, …, *v*_*k*_, *u*_1_, …, *u*_*l*_*}*. The negative samples are then

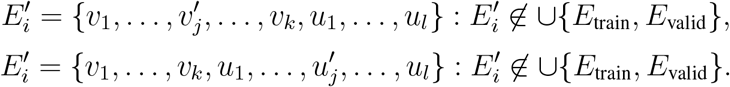

Furthermore, for hypergraphs with fixed-length hyperedges like the polypharmacy dataset [36], we propose an enhanced negative sampling strategy. Let the fixed length *k* = 3 and *E*_*i*_ = *{v*_1_, *v*_2_, *v*_3_*}*, we generate the negative sample as

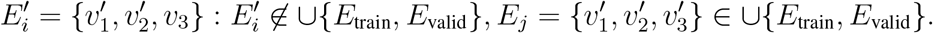

so that the altered nodes in the negative sample are guaranteed to interact in another hyperedge.

##### Attention Network

As in Figure 5c, the attention network follows the design in [28] with weight matrices 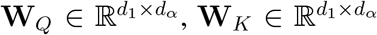, and 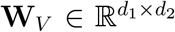. Given the a node set *E* of size k and the node embeddings 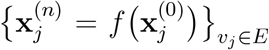, the normalized attention coeffcient *α*_*il*_ regarding node *v*_*i*_ and node *v*_*l*_ is computed as

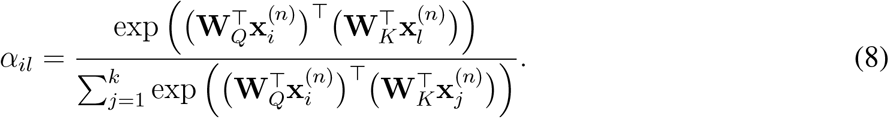

Then, the dynamic embedding of each node *v*_*i*_ *∈ E* is computed as

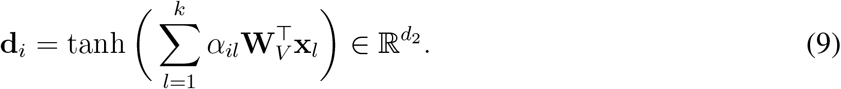

And with weight matrix 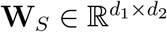, the static embedding of each node *v*_*i*_ *∈ E* is computed as

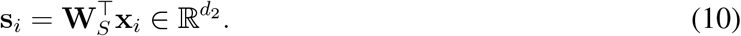

Finally, with another layer 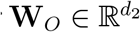, the output of the attention network is then the probability that the node set is a hyperedge:

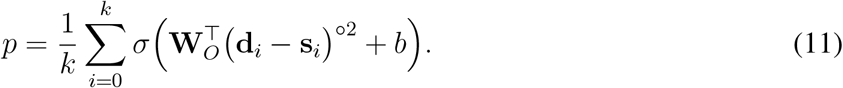

where *σ* is the sigmoid function and *b ∈* R. Notation ∘ denotes element-wise power operation.

#### Downstream Tasks

We evaluate how well the model learns node representation through node-level downstream tasks. Loading the saved base model and adding a fully connected layer for prediction, we fine-tune the model by the downstream task. Based on the pre-trained model *f* (*·*) that generates node embeddings with the information it leveraged from the pre-training task, we can utilize it to do node-level downstream tasks in a wide range of applications.

Let *h*(*·*) denote the fully connected layer that is task-specific for different data. For node classification tasks with *C* classes as in Section, we have 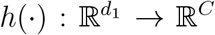. Given the training set *S*_train_ with input feature 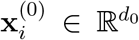 for each node *i* the label *y*_*i*_ *∈* [*C*] representing the node class, we minimize the following cross-entropy loss with regard to each training batch *S*:

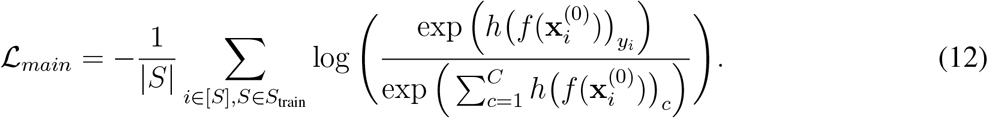

For continual signal prediction tasks as in Section, we have 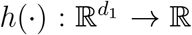. Given the training set *S*_train_ with input feature 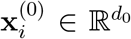 for each node *i* the label *y*_*i*_ *∈* R, we minimize the following mean squared error with regard to each training batch *S*:

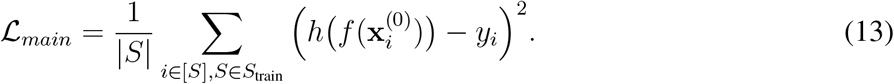

In the downstream tasks, we keep fine-tuning the base convolutional network *f* (*·*) with the task-specific fully connected layer *h*(*·*).

## Acknowledgements

This work was supported in part by the National Institutes of Health Common Fund 4D Nucleome Program grant UM1HG011593 (J.M.), National Institutes of Health Common Fund Cellular Senescence Network Program grant UG3CA268202 (J.M.), National Institutes of Health grants R01HG007352 (J.M.) and R01HG012303 (J.M.), and National Science Foundation grants 1855099 (Q.G.), 1903202 (Q.G.), and 2247426 (Q.G.). J.M. was additionally supported by a Guggenheim Fellowship from the John Simon Guggenheim Memorial Foundation.

## Author Contributions

Conceptualization, Y.D., R.Z., J.M., and Q.G.; Software, Y.D..; Investigation, Y.D., R.Z., P.X., J.M., and Q.G.; Writing, Y.D., R.Z., P.X., J.M., and Q.G.; Funding Acquisition, J.M. and Q.G.

## Competing Interests

The authors declare no competing interests.

## Notes

### Competing Interest Statement

The authors have declared no competing interest.

## References

[1] Agarwal, S. et al. Beyond pairwise clustering. In 2005 IEEE Computer Society Conference on Computer Vision and Pattern Recognition (CVPR’05), vol. 2, 838–845 (IEEE, 2005).

[2] Agarwal, S., Branson, K. & Belongie, S. Higher order learning with graphs. In Proceedings of the 23rd international conference on Machine learning, 17–24 (2006).

[3] Satchidanand, S. N., Ananthapadmanaban, H. & Ravindran, B. Extended discriminative random walk: a hypergraph approach to multi-view multi-relational transductive learning. In Twenty-Fourth International Joint Conference on Artificial Intelligence (2015).

[4] Sun, L., Ji, S. & Ye, J. Hypergraph spectral learning for multi-label classification. In KDD (2008).

[5] Yadati, N. et al./person-group>. Hypergcn: A new method for training graph convolutional networks on hyper-graphs. In Wallach, H.et al. (eds.) Advances in Neural Information Processing Systems, vol. 32 (Curran Associates, Inc., 2019).

[6] Dong, Y., Sawin, W. & Bengio, Y. Hnhn: Hypergraph networks with hyperedge neurons. arXiv preprint arXiv:2006.12278 (2020).

[7] Arya, D., Gupta, D. K., Rudinac, S. & Worring, M. Hypersage: Generalizing inductive representation learning on hypergraphs. arXiv preprint arXiv:2010.04558 (2020).

[8] Bai, S., Zhang, F. & Torr, P. H. Hypergraph convolution and hypergraph attention. Pattern Recognition 110, 107637 (2021).

[9] Yi, J. & Park, J. Hypergraph convolutional recurrent neural network. In Proceedings of the 26th ACM SIGKDD International Conference on Knowledge Discovery & Data Mining, 3366–3376 (2020).

[10] Sun, X. et al. Heterogeneous hypergraph embedding for graph classification (2021).

[11] Doersch, C., Gupta, A. & Efros, A. A. Unsupervised visual representation learning by context prediction. In Proceedings of the IEEE international conference on computer vision, 1422–1430 (2015).

[12] Kolesnikov, A., Zhai, X. & Beyer, L. Revisiting self-supervised visual representation learning. In Proceedings of the IEEE/CVF conference on computer vision and pattern recognition, 1920–1929 (2019).

[13] Girshick, R., Donahue, J., Darrell, T. & Malik, J. Rich feature hierarchies for accurate object detection and semantic segmentation. In Proceedings of the IEEE conference on computer vision and pattern recognition, 580–587 (2014).

[14] He, K., Girshick, R. & Dollar, P. Rethinking imagenet pre-training. In Proceedings of the IEEE/CVF International Conference on Computer Vision (ICCV) (2019).

[15] Devlin, J., Chang, M.-W., Lee, K. & Toutanova, K. BERT: Pre-training of deep bidirectional transformers for language understanding. In Proceedings of the 2019 Conference of the North American Chapter of the Association for Computational Linguistics: Human Language Technologies, Volume 1 (Long and Short Papers), 4171–4186 (Association for Computational Linguistics, Minneapolis, Minnesota, 2019).

[16] Hu, W. et al. Strategies for pre-training graph neural networks. In International Conference on Learning Representations (ICLR) (2020).

[17] You, Y., Chen, T., Wang, Z. & Shen, Y. When does self-supervision help graph convolutional networks? In International Conference on Machine Learning, 10871–10880 (PMLR, 2020).

[18] Hu, Z., Dong, Y., Wang, K., Chang, K.-W. & Sun, Y. Gpt-gnn: Generative pre-training of graph neural networks. In Proceedings of the 26th ACM SIGKDD Conference on Knowledge Discovery and Data Mining (2020).

[19] Wu, J. et al. Self-supervised graph learning for recommendation. In Proceedings of the 44th international ACM SIGIR conference on research and development in information retrieval, 726–735 (2021).

[20] Jin, W. et al. Automated self-supervised learning for graphs. In International Conference on Learning Representations (2022).

[21] Hwang, D. et al./person-group>. Self-supervised auxiliary learning with meta-paths for heterogeneous graphs. In Larochelle, H., Ranzato, M., Hadsell, R., Balcan, M. F. & Lin, H. (eds.) Advances in Neural Information Processing Systems, vol. 33, 10294–10305 (Curran Associates, Inc., 2020).

[22] Hao, B., Zhang, J., Yin, H., Li, C. & Chen, H. Pre-training graph neural networks for cold-start users and items representation. In Proceedings of the 14th ACM International Conference on Web Search and Data Mining, 265–273 (2021).

[23] Sun, K., Lin, Z. & Zhu, Z. Multi-stage self-supervised learning for graph convolutional networks on graphs with few labeled nodes. Proceedings of the AAAI Conference on Artificial Intelligence 34, 5892–5899 (2020).

[24] Xia, X. et al. Self-supervised hypergraph convolutional networks for session-based recommendation. In Proceedings of the AAAI Conference on Artificial Intelligence, vol. 35, 4503–4511 (2021).

[25] Yu, J. et al. Self-supervised multi-channel hypergraph convolutional network for social recommendation. In Proceedings of the Web Conference 2021, 413–424 (2021).

[26] Xia, L., Huang, C. & Zhang, C. Self-supervised hypergraph transformer for recommender systems. In Proceedings of the 28th ACM SIGKDD Conference on Knowledge Discovery and Data Mining, 2100–2109 (2022).

[27] Du, B., Yuan, C., Barton, R., Neiman, T. & Tong, H. Hypergraph pre-training with graph neural networks. arXiv preprint arXiv:2105.10862 (2021).

[28] Zhang, R., Zou, Y. & Ma, J. Hyper-SAGNN: a self-attention based graph neural network for hypergraphs. In International Conference on Learning Representations (2020).

[29] Tu, K., Cui, P., Wang, X., Wang, F. & Zhu, W. Structural deep embedding for hyper-networks. In AAAI (2018).

[30] Chen, D. et al. Measuring and relieving the over-smoothing problem for graph neural networks from the topological view. In Proceedings of the AAAI Conference on Artificial Intelligence, vol. 34, 3438–3445 (2020).

[31] Rong, Y., Huang, W., Xu, T. & Huang, J. Dropedge: Towards deep graph convolutional networks on node classification. In International Conference on Learning Representations (2020).

[32] Kipf, T. N. & Welling, M. Semi-supervised classification with graph convolutional networks. In International Conference on Learning Representations (ICLR) (2017).

[33] Quinodoz, S. A. et al. Higher-order inter-chromosomal hubs shape 3d genome organization in the nucleus. Cell 174, 744–757 (2018).

[34] Zhang, R. & Ma, J. Matcha: Probing multi-way chromatin interaction with hypergraph representation learning. Cell Systems 10, 397–407 (2020).

[35] Lieberman-Aiden, E. et al. Comprehensive mapping of long-range interactions reveals folding principles of the human genome. Science 326, 289–293 (2009).

[36] Zitnik, M., Agrawal, M. & Leskovec, J. Modeling polypharmacy side effects with graph convolutional networks. Bioinformatics 34, i457–i466 (2018).

[37] Nováček, V. & Mohamed, S. K. Predicting polypharmacy side-effects using knowledge graph embeddings. AMIA Summits on Translational Science Proceedings 2020, 449 (2020).

[38] Xu, K., Hu, W., Leskovec, J. & Jegelka, S. How powerful are graph neural networks? In International Conference on Learning Representations (2019).

[39] Feng, Y., You, H., Zhang, Z., Ji, R. & Gao, Y. Hypergraph neural networks. In Proceedings of the AAAI Conference on Artificial Intelligence, vol. 33, 3558–3565 (2019).

[40] Wu, Z. et al. A comprehensive survey on graph neural networks. IEEE transactions on neural networks and learning systems 32, 4–24 (2020).

[41] Li, G., Müller, M., Thabet, A. & Ghanem, B. Deepgcns: Can gcns go as deep as cnns? In The IEEE International Conference on Computer Vision (ICCV) (2019).

[42] Chen, G. & Zhang, J. Preventing over-smoothing for hypergraph neural networks. arXiv preprint arXiv:2203.17159 (2022).

