## Supplemental Information for "PhyGCN: Pre-trained Hypergraph Convolutional Neural Networks with Self-supervised Learning"

#### A Related Work

Interactions naturally occur among objects in real-world datasets. The pairwise interactions are conveniently modeled as graphs, facilitating their analysis through deep learning models such as graph neural networks (GNNs) [1–4]. GNNs leverage the graph structure to generate representations of unknown nodes for various downstream tasks such as node classification [5, 6], predicting possible unknown interactions between two nodes [7], and classification of graphs [8]. However, real-world data often exhibit relationships that are more complex than pairwise interactions; higher-order interactions involve more than two nodes. For example, in the Cora co-citation network [9], multiple papers are connected by citations from a common paper. Similarly, in the drug side-effect network [10], interactions may exist among multiple drugs and side effects. Such graph structures with higher-order interactions defines as **hypergraph** [11], extending the concept of conventional graphs with pairwise interactions. In a hypergraph, these interactions are termed as hyperedges. Being a more complex data structure, hypergraphs contain richer information comapred to graphs with only pairwise edges. However, this complexity poses challenges to classical GNN models, which requires the development of new methods to represent and learn the interactions among arbitrarily many nodes instead of just pairwise.

Reflecting on the well-explored domains GNNs, a mainstream approach for utilizing graph structures is Graph Convolutional Networks (GCNs) [2]. TGCNs aim to learn node representations by aggregating information from neighboring nodes and combining it with the node's own information. A detailed illustration of GCNs is **Methods**. While GCNs have proven effective in representation learning for graphs, the difference in structures between graphs and hypergraphs poses a challenge when adapting GCNs to hypergraphs, especially in capturing the additional information brought by the hyperedges. A straightforward approach to tackle this challenge is to transform the hypergraph into a conventional pairwise graph through clique expansion [12], then applying standard GCN methods. Essentially, this entails treating each pair of nodes within a hyperedge as an edge in conventional graphs. Analogous to such expansion, simple convolutional architectures for hypergraphs have been proposed and used in several works [13, 14]. [13] is the first work to introduce a simple neural network structure with a convolutional layer for hyperedges.

Several other works aim to enhance the performance of hypergraph GCNs by altering the aggregation and combination of neighboring node information. Technically, this variety stems from different designs of the hypergraph's Laplacian matrix (elaborated in **Methods**). [15] also approximates hyperedges with a set of pairwise edges, proposing the HyperGCN scheme that achieves the best result and FastHyperGCN that trains faster albeit with compromised performance. Subsequently, to better capture hyperedge information, [16] proposes Hypergraph Networks with Hyperedge Neurons (HNHN), a hypergraph convolutional network applying nonlinear activation functions to both nodes and hyperedges. This model computes embeddings for nodes and hyperedges. However, under varied train-test data splits on multiple hypergraph data, these methods could exhibit instability and perform worse than a common GCN learning on hypergraphs expanded as regular graphs. Other attempts to adapt graph architectures to hypergraphs either achieve similar or slightly improved performance compared to HyperGCN [14, 17, 18] or are evaluated under different conditions such as a larger amount of training data [19].

Expansions like clique expansion assume a hyperedge's decomposability, implying that any subset of nodes in a hyperedge can form another hyperedge. However, this assumption becomes unreasonable when a hyperedge contains nodes of different types. For instance, a hyperedge may represent a triplet relationship between a user, drug, and reaction. Treating such a hyperedge as pairwise edges could be detrimental to downstream tasks. This issue is similarly noted in [20], where the authors attempt to model the hypergraph with a simple encoder-decoder model. Yes, as evidenced in our empirical analysis for the multiway chromatin interaction data, such a model does not match the capability of a deep hypergraphgraph convolutional model.

While relatively few works explore self-supervised training or pre-training on hypergraphs, the field of graphs has seen many mature works. Comprehensive studies [21–24] have introduced and examined multiple self-supervised tasks for graphs at both the node and graph level. Additionally, some works [25–28] are tailored for specific settings, employing one or more similar self-supervised tasks to enhance model performance and adapt to chosen scenarios. In another subfield of self-supervised training, contrastive learning for GNNs has also been explored in recent studies [29–31].

Self-supervised learning and pre-training for hypergraphs remain largely unexplored. The majority of works [32–34] focus on specific applications in recommender systems. [32] introduces Dual Channel Hypergraph Convolutional Networks (DHCN), explicitly designed for session-based recommendation. DHCN's self-supervised learning task seeks to maximize mutual information between session representations learned by a graph convolutional network and DHCN, aiming to enhance hypergraph network learning. [33] also focuses on recommender systems, leveraging higher-order interactions to improve recommendation quality. They introduce a self-supervised task to compensate for the loss incured by their multi-channel hypergraph convolutional network, aiming to maximize mutual information concerning the representations of the user node, the neighboring sub-hypergraph of the user, and the entire hypergraph. [34] aims to enhance recommendation and user representations via a hypergraph transformer network that uses a cross-view generative self-supervised learning for data augmentation. Lastly, [35] proposes pre-training with GNNs on hypergraphs for hyperedge classification. This method uses GNNs with self-supervised pre-training tasks, achieving state-of-the-art performances on both inductive and transductive hyperedge classification.

### B Experimental Details

#### B.1 Model Parameter Setting in Pre-training

For PhyGCN, we consider three hypergraph convolutional layers, each with a uniform hidden layer size of 128 and an output embedding size of 64. In the multi-head attention layer used for pre-training, we set the number of heads to 8, aligning with setting in [36]. The number of training epochs is set to be 300, and the training batch of the pre-training task contains 32 positive samples while we generate 5 negative samples with regard to each positive sample. We use Adam optimizer of learning rate 0.001 with weight decay 5e-4. The dropout rate is set to 0.5, and the coefficient for dropedge is set to 0.3: masking 30% of the values of the adjacency matrix $\widehat{\mathbf{A}}$ at each training iteration.

#### B.2 Supplementary Information for the Citation Networks Data

**Baselines.** We compare our approach with two recent baselines on hypergraph neural networks alongside an additional baseline derived from modifying the graph convolutional network:

- HyperGCN, proposed by [15], represents their best performing model. This method incorporates node features when calculating the Laplacian to improve performances. However, such computation significantly increases the computation cost for the method.
- HNHN, introduced by [16], divides the aggregation process into two steps, computing embeddings for hyperedges and nodes separately.
- GCN, here we expand the hypergraph into graphs with pairwise edges through clique expansion and apply a straightforward GCN model [2] for prediction.

**Datasets.** The details of the datasets used for the evaluation of node classification are shown in **Table** S1. It is noteworthy that the data used by [15] contains a lot of isolated nodes, implying that a significant portion of nodes, not part of any hyperedge, are classified solely based on their feature information. This is not an ideal setting for evaluating the capacity of a graph/hypergraph neural network in capturing structural information. We opt for a similar setting to [16], where we remove the isolated nodes from the raw data. Moreover, since the compared works [15, 16] use different train-test ratios for each dataset, we make a uniform setting of ratios for all data and models: 0.5%, 1%, 2%, 4%. Specifically, we conduct four experiments on each dataset for each model, incrementally increasing the training data ratio from 0.5% to 4%. Training data with a larger proportion includes the nodes from the training data of a smaller proportion. Lastly, we perform 10 random splits for each ratio of each dataset to evaluate the average performance alongside the standard deviation.

**Training.** All baselines are trained using the Adam optimizer [37] as specified in their respective papers. The selected hyperparameters for each baseline also adhere to those suggested in the respective papers. For PhyGCN, we use SGD of learning rate 0.003 with momentum 0.9 for the node classification task.

| | DBLP (co-authorship) | Pubmed (co-citation) | Cora (co-authorship) | Cora (co-citation) | Citeseer (co-citation) |
|---|---|---|---|---|---|
| # nodes, $\|V\|$ | 41302 | 3840 | 2388 | 1434 | 1458 |
| # hyperedges, $\|E\|$ | 22363 | 7963 | 1072 | 1579 | 1079 |
| avg hyperedge size | 4.5 ± 5.4 | 4.3 ± 5.7 | 4.3 ± 4.2 | 3.0 ± 1.0 | 3.2 ± 2.0 |
| # features, d | 1425 | 500 | 1433 | 1433 | 3703 |
| # classes, q | 6 | 3 | 7 | 7 | 6 |

**Table S1:** The statistics for the citation dataset used in node classification, where each data has the hyperedge either as a co-authorship relation (papers written by the same author) or as a co-citation relation (papers cited by the same paper).

**Detailed Empirical Results** Detailed results of the average test accuracy $\pm$ standard deviation for all models on all datasets are shown in **Table** S2. We found that PhyGCN outperforms all baselines on each dataset regardless of the training data ratios. On smaller hypergraphs like Cora and Citeseer, PhyGCN is able to leverage more information with self-supervised learning and outperforms all baselines by a large margin. Similarly, in scenarios with low training data ratio on large networks, PhyGCN maintains remarkable robustness and exemplary performance, while other hypergraph methods such as HyperGCN and HNHN falter. An ablation study of the pre-training scheme presents results of our hypergraph convolutional network both with and without pre-training. A direct comparison shows the substantial benefit of pre-training. For instance, on the Cora dataset, where the plain model underperforms when compared to the vanilla GCN, pre-training enables the hypergraph convolutional network to better capture higher-order information, and significantly outperforming GCN.

#### B.3 Supplementary Information for the Multiway Chromatin Interaction Data

**Baselines.** For the hyperedge prediction task on multi-way chromatin interaction, we evaluated against a recent work, MATCHA [38], which employs hypergraph representation learning techniques on a specially constructed hypergraph from multi-way chromatin interaction data, aimed at denoising the data and making *de-novo* predictions of yet undetected multi-way chromatin interactions. For the baseline of **Fig. 3c** and **Fig. 3d**, we used A/B compartment scores [39], a 3D genome features that is known to correlate with DNA replication timing. The A/B compartment scores are calculated using the Cooler software [40] by taking the SPRITE contact map as input.

**Dataset.** To systematically evaluate if our proposed new approach could improve the analysis of multi-way chromatin interaction data, we use the SPRITE data [41] from the GM12878 lymphoblastoid human cell line following the same processing procedure in MATCHA [38]. Specifically, genomic bins at the 1Mb resolution are considered as nodes, and interactions involving multiple loci are treated as hyperedges. In **Fig. 3a**, a split by chromosome indices is performed, where nodes odd chromosome indices constitute the training data, and those from even indices form the test data. For cross-validation, a swapping of the training and test set is conducted, followed by the reporting of the average accuracy for each method. **Fig. 3b** displays a comparison against MATCHA [38] under a more challenging training split, constructing the hypergraph from SPRITE data with 20% designated as training set and the remaining

| Dataset | Model | Training Ratio | | | |
|---|---|---|---|---|---|
| | | 0.5% | 1% | 2% | 4% |
| Citeseer | HyperGCN | 30.71 ± 7.0 | 41.02 ± 10.6 | 51.50 ± 8.5 | 63.10 ± 4.6 |
| | HNHN | 26.33 ± 7.5 | 28.61 ± 5.9 | 34.65 ± 6.9 | 45.35 ± 6.5 |
| | GCN | 32.66 ± 10.4 | 44.02 ± 9.7 | 53.91 ± 6.4 | 62.54 ± 3.7 |
| | Our model (plain) | 32.74 ± 6.2 | 38.99 ± 6.5 | 49.21 ± 0.5 | 56.72 ± 3.2 |
| | Our model (pre-trained) | **47.35 ± 8.4** | **54.22 ± 7.2** | **62.87 ± 4.3** | **67.02 ± 1.0** |
| Cora (co-authorship) | HyperGCN | 33.08 ± 8.6 | 39.50 ± 4.7 | 49.24 ± 4.8 | 57.36 ± 5.2 |
| | HNHN | 30.48 ± 5.5 | 38.23 ± 10.4 | 53.53 ± 4.0 | 63.51 ± 3.1 |
| | GCN | 38.92 ± 9.3 | 54.34 ± 7.4 | 60.35 ± 5.3 | 69.00 ± 2.2 |
| | Our model (plain) | 38.21 ± 7.6 | 50.21 ± 5.6 | 59.66 ± 5.7 | 68.53 ± 2.6 |
| | Our model (pre-trained) | **49.43 ± 6.5** | **60.75 ± 5.8** | **67.21 ± 3.6** | **71.86 ± 1.9** |
| Cora (co-citation) | HyperGCN | 33.06 ± 6.2 | 37.31 ± 7.3 | 45.97 ± 6.9 | 56.72 ± 5.6 |
| | HNHN | 26.16 ± 4.5 | 28.44 ± 4.7 | 38.51 ± 7.5 | 55.60 ± 3.3 |
| | GCN | 39.52 ± 9.7 | 48.37 ± 6.8 | 58.72 ± 7.0 | 71.15 ± 3.5 |
| | Our model (plain) | 33.37 ± 6.6 | 38.31 ± 5.6 | 48.77 ± 5.5 | 62.28 ± 3.2 |
| | Our model (pre-trained) | **48.82 ± 8.2** | **58.46 ± 5.5** | **65.71 ± 4.5** | **72.35 ± 3.2** |
| DBLP | HyperGCN | 65.35 ± 4.3 | 66.30 ± 6.2 | 63.79 ± 4.7 | 65.84 ± 6.4 |
| | HNHN | 74.41 ± 1.9 | 79.86 ± 0.9 | 82.19 ± 0.4 | 83.73 ± 0.2 |
| | GCN | 81.11 ± 2.9 | 83.65 ± 2.0 | 85.32 ± 0.7 | 85.84 ± 0.5 |
| | Our model (plain) | 81.90 ± 0.6 | 83.71 ± 0.4 | 85.24 ± 0.2 | 86.52 ± 0.2 |
| | Our model (pre-trained) | **84.69 ± 0.6** | **86.12 ± 0.4** | **86.96 ± 0.2** | **87.80 ± 0.2** |
| PubMed | HyperGCN | 67.84 ± 6.3 | 72.29 ± 5.4 | 77.10 ± 2.1 | 79.63 ± 1.0 |
| | HNHN | 63.61 ± 2.9 | 68.86 ± 3.9 | 72.24 ± 3.2 | 76.49 ± 1.6 |
| | GCN | 70.29 ± 4.4 | 74.31 ± 2.8 | 77.92 ± 1.5 | 79.66 ± 1.0 |
| | Our model (plain) | 70.96 ± 3.7 | 75.07 ± 2.2 | 78.01 ± 1.0 | 79.84 ± 0.5 |
| | Our model (pre-trained) | **74.66 ± 3.9** | **78.00 ± 1.6** | **79.09 ± 1.6** | **79.96 ± 0.9** |

**Table S2:** Test accuracy for different datasets at different data split ratio (training data % = 0.5% , 1% , 2% , 4%). We report mean test accuracy ± standard deviation among $10$ random train-test split for each split ratio.

| | Accuracy |
|---|---|
| Plain | 28.42 % |
| Pre-trained | **80.79 %** |

**Table S3:** Classification on GM12878.

| | MATCHA | Our Method |
|---|---|---|
| Size= 2 | (0.898, 0.787) | **(0.934, 0.827)** |
| Size= 3 | (0.988, 0.962) | **(0.996, 0.977)** |
| Size= 4 | (0.996, 0.981) | **(0.998, 0.992)** |

**Table S4:** Hyperedge prediction on SPRITE data. Size denotes the length of the hyperedge. We report the (AUROC, AUPR) scores.

80% as test data. **Fig. 3c** and **Fig. 3d**, the replication timing for each genomic bin is quantified using Repli-seq [42].

**Detailed Empirical Results.** **Table** S3 shows the detailed results for **Fig. 3a**. **Table** S4 shows the detailed results for **Fig. 3b**.

##### B.4 Supplementary Information for Polypharmacy Side Effect Data

**Baselines.** We compared our approach with several methods on polypharmacy side effect prediction:

- Decagon, proposed by [10] as the initial work that studies the polypharmacy side-effect data.
- ComplEX, proposed in [43] that has the best reported performances.
- HyperSAGNN, with the encoder-decoder model for representation learning as proposed in [36].

**Dataset.** We use the polypharmacy side-effect dataset [10], which contains 645 unique drugs and 63,473 drug-drug interactions. Within the hypergraph framework, there are 4,651,131 hyperedges representing the drug-drug side-effect associations, featuring 1,317 unique side effects. The evaluation on side-effect prediction was conducted across four distinct train-test split scenarios:

- *Original*: The same as in [10], where the test set only contains side effects associated with more than 500 drug combinations, whilst side-effects liked with less than 500 drug combinations are allocated to the training set.
- *Random*: All side-effects are utilized for both training and testing, distributed randomly across training and test sets.
- *Expected*: A subset of 30 side-effects, already associated with a single drug, were deemed as expected, leading to 20,787 expected drug-drug side-effect associations. Despite adhering to a random split for training and test data, the "expected" scenario only tests on the expected triplets within the test set.

| Model and training scheme | Accuracy |
|---|---|
| Plain | 21.49 % |
| With pre-training, random negative sampling | 33.65 % |
| With pre-training, modified negative sampling | **35.23 %** |

**Table S5:** Classification of side-effects.

| | Decagon | ComplEX | Hyper-SAGNN | Our Method |
|---|---|---|---|---|
| Original | (0.814, 0.737) | (0.929, 0.914) | (0.938, 0.921) | **(0.945, 0.931)** |
| Random | (0.801, 0.729) | (0.920, 0.903) | (0.933, 0.912) | **(0.943, 0.926)** |
| Expected | (0.807, 0.733) | (0.943, 0.931) | (0.938, 0.924) | **(0.947, 0.932)** |
| Unexpected | (0.786, 0.713) | (0.917, 0.890) | (0.935, 0.918) | **(0.944, 0.922)** |

**Table S6:** Performances on different data split. Random negative sampling. We report the (AUROC, AUPR) scores for each model on each data split.

- *Unexpected*: Similarly, whilst maintaining a random split for training and test data, testing is conducted solely on the unexpected triplets in the test set.

For the side effect categorization task, we consider side effects with known categories from [10], yielding 561 side effects with 37 categories.

**Detailed Empirical Results.** In **Table** S6 and **Table** S7, we show detailed results of the AUROC and AUPR scores for all models on the four different evaluation sets. The AUROC scores in **Table** S6 correspond to **Fig.** **3e** and the AUROC scores in **Table** S7 correspond to **Fig.** **3f**. The AUPR scores of **Table** S6 showed a similar pattern as AUROC scores, where hypergraph methods tend to maintain their performance on expected and unexpected test data while graph methods exhibit a decline in performance. Similarly, for the AUPR scores of **Table** S7, the enhanced negative sampling strategy brings more challenge to the task and forces the networks to learn the more subtle differences between positive and negative samples.

Our results demonstrated the adeptness of our model in learning higher-order interactions effectively. We also studied how learning within the hypergraph structure could improve our model's performance on tasks pertinent to the nodes (drugs, side effects). We used the side effect category information from [10], where a proportion of the side effects within the network were classified into high-level categories. As shown in **Table** S5, with pre-training on the hypergraph structure, our model exhibited enhanced categorization capabilities, even in the absence of additional information. Furthermore, the performance metric could be improved through the implementation of a better negative sampling strategy.

| | ComplEX | Hyper-SAGNN | Our Method |
|---|---|---|---|
| Original | (0.724, 0.702) | (0.716, 0.702) | **(0.744, 0.731)** |
| Random | (0.723, 0.699) | (0.715, 0.703) | **(0.743, 0.730)** |
| Expected | (0.739, 0.722) | (0.719, 0.709) | **(0.746, 0.732)** |
| Unexpected | (0.703, 0.686) | (0.718, 0.703) | **(0.742, 0.729)** |

**Table S7:** Performances on different data split. Modified negative sampling. We report the (AUROC, AUPR) scores for each model on each data split.

| | GPS | MovieLens | drug | wordnet |
|---|---|---|---|---|
| Encoder | (0.930, 0.744) | (0.926, 0.793) | (0.961, 0.888) | (0.890, 0.694) |
| Conv | **(0.941, 0.738)** | **(0.938, 0.801)** | (0.962, 0.890) | **(0.896, 0.710)** |

**Table S8:** The experiment results for the hyperedge prediction task over the four heterogeneous hypergraph data with uniform-length hyperedges. We report (AUROC, AUPR) to evaluate how well our model learns the structural knowledge. Our convolutional model is compared with the encoder-decoder model used in [36], both using the attention layer for prediction.

##### B.5 The Base Convolutional Model Learns Hyperedge Information Well

We further evaluated how well the hypergraph convolutional network in PhyGCN captures the hyperedge information. We conducted evaluation on the the four datasets used in the most current works on hyperedge prediction [20, 36]. These datasets, shown below, have uniform-length hyperedges and no node features or node labels.

- GPS [44]: hyperedges represent (user, location, activity) relations.
- MovieLens [45]: hyperedges represent (user, movie, tag) relations.
- drug: hyperedges represent (user, drug, reaction) relations.
- wordnet [46]: hyperedges represent (head entity, relation, tail entity) relations within words.

Hyperedge prediction methods need to use structural information as the features to make predictions. Moreover, these datasets are heterogeneous, meaning that the node types within a hyperedge are different and therefore one could not simply expand the hyperedge into pairwise edges. Details of the experiment results are reported in **Table** S8. We evaluated our base convolutional model against the encoder-decoder model used in [20, 36] for representation learning, using the same attention network from [36] for combining the node embeddings and making the prediction. The performance is evaluated by the AUROC and the AUPR score. We found that, with the introduction of convolutional architecture, our model performs either better than or on par with the encoder-decoder model. We note that the performances for the four datasets are already very high with the encoder-decoder model, our base model still exhibit enhanced capability in capturing the hyperedge information.

In **Table** S9, we extended our evaluation to five datasets for the main task of node classification. These datasets are citation networks where each node represents a publication and each hyperedge represents either co-citation or co-authorship relation. These data are different from the above setting, where the hypergraph is homogeneous with variable hyperedge size. We note that, for the previous hyperedge prediction data, the methods typically use features constructed from the hypergraph structure: a detailed formulation of the construction of input features are presented in **Methods**. However, as these five datasets have node features themselves, we additionally add two modified baselines of the encoder-decoder model for a more comprehensive evaluation. Specifically, we compare with encoder-decoder that takes in the constructed features, the node features, and concatenation of the two. Our model soley utilizes the node feature for these five datasets. We found that our convolutional base model outperforms the three baselines on AUROC score consistently across the five datasets. Such results underscore the

| | Citeseer | Cora (co-authorship) | Cora (co-cocitation) | DBLP | PubMed |
|---|---|---|---|---|---|
| Encoder-1 | (0.804, 0.573) | (0.827, 0.626) | (0.809, 0.591) | (0.825, 0.655) | (0.906, 0.722) |
| Encoder-2 | (0.833, 0.612) | (0.798, 0.553) | (0.849, 0.590) | (0.920, 0.727) | (0.919, 0.730) |
| Encoder-3 | (0.803, 0.583) | (0.826, 0.660) | (0.839, 0.607) | **(0.930, 0.773)** | **(0.920, 0.752)** |
| Conv | **(0.866, 0.605)** | **(0.861, 0.657)** | **(0.861, 0.597)** | (0.923, 0.755) | **(0.926, 0.727)** |

**Table S9:** The experiment results for hyperedge prediction over the five homogeneous datasets with variable-length hyperedges. We report (AUROC, AUPR) similarly. We compare our convolutional model with the encoder-decoder model that takes in three different features. Encoder-1: original features computed as in [36]. Encoder-2: node features provided by the data. Encoder-3: concatenation of the two.

potential of our base model to effectively and efficiently utilize both the node features and the structural information of the hypergraph. For the hyperedge prediction task only, the base convolutional model has a stably better performance compared to the simple model used in previous works. Such evaluation confirms that the model learns the structural knowledge well in this pre-training task.

#### References


[1] Li, Y., Zemel, R., Brockschmidt, M. & Tarlow, D. Gated graph sequence neural networks. In *Proceedings of ICLR'16* (2016).

[2] Kipf, T. N. & Welling, M. Semi-supervised classification with graph convolutional networks. In *International Conference on Learning Representations (ICLR)* (2017).

[3] Veličković, P. *et al.* Graph Attention Networks. *International Conference on Learning Representations* (2018). Accepted as poster.

[4] Xu, K., Hu, W., Leskovec, J. & Jegelka, S. How powerful are graph neural networks? In *International Conference on Learning Representations* (2019).

[5] Yao, L., Mao, C. & Luo, Y. Graph convolutional networks for text classification. In *Proceedings of the AAAI conference on artificial intelligence*, vol. 33, 7370–7377 (2019).

[6] Hong, D. *et al.* Graph convolutional networks for hyperspectral image classification. *IEEE Transactions on Geoscience and Remote Sensing* **59**, 5966–5978 (2020).

[7] Yuan, Y. & Bar-Joseph, Z. Gcng: graph convolutional networks for inferring gene interaction from spatial transcriptomics data. *Genome biology* **21**, 1–16 (2020).

[8] Zhang, M., Cui, Z., Neumann, M. & Chen, Y. An end-to-end deep learning architecture for graph classification. In *Proceedings of the AAAI conference on artificial intelligence*, vol. 32 (2018).

[9] McCallum, A. K., Nigam, K., Rennie, J. & Seymore, K. Automating the construction of internet portals with machine learning. *Information Retrieval* **3**, 127–163 (2000).

[10] Zitnik, M., Agrawal, M. & Leskovec, J. Modeling polypharmacy side effects with graph convolutional networks. *Bioinformatics* **34**, i457–i466 (2018).

[11] Bretto, A. Hypergraph theory. *An introduction. Mathematical Engineering. Cham: Springer* (2013).

[12] Sun, L., Ji, S. & Ye, J. Hypergraph spectral learning for multi-label classification. In *KDD* (2008).

[13] Feng, Y., You, H., Zhang, Z., Ji, R. & Gao, Y. Hypergraph neural networks. In *Proceedings of the AAAI Conference on Artificial Intelligence*, vol. 33, 3558–3565 (2019).

[14] Bai, S., Zhang, F. & Torr, P. H. Hypergraph convolution and hypergraph attention. *Pattern Recognition* **110**, 107637 (2021).

[15] Yadati, N. *et al.* Hypergcn: A new method for training graph convolutional networks on hypergraphs. In Wallach, H. *et al.* (eds.) *Advances in Neural Information Processing Systems*, vol. 32 (Curran Associates, Inc., 2019).

[16] Dong, Y., Sawin, W. & Bengio, Y. Hnhn: Hypergraph networks with hyperedge neurons. *arXiv preprint arXiv:2006.12278* (2020).

[17] Arya, D., Gupta, D. K., Rudinac, S. & Worring, M. Hypersage: Generalizing inductive representation learning on hypergraphs. *arXiv preprint arXiv:2010.04558* (2020).

[18] Yi, J. & Park, J. Hypergraph convolutional recurrent neural network. In *Proceedings of the 26th ACM SIGKDD International Conference on Knowledge Discovery & Data Mining*, 3366–3376 (2020).

[19] Sun, X. *et al.* Heterogeneous hypergraph embedding for graph classification (2021).

[20] Tu, K., Cui, P., Wang, X., Wang, F. & Zhu, W. Structural deep embedding for hyper-networks. In *AAAI* (2018).

[21] Hu, W. *et al.* Strategies for pre-training graph neural networks. In *International Conference on Learning Representations (ICLR)* (2020).

[22] You, Y., Chen, T., Wang, Z. & Shen, Y. When does self-supervision help graph convolutional networks? In *International Conference on Machine Learning*, 10871–10880 (PMLR, 2020).

[23] Wu, J. *et al.* Self-supervised graph learning for recommendation. In *Proceedings of the 44th international ACM SIGIR conference on research and development in information retrieval*, 726–735 (2021).

[24] Jin, W. *et al.* Automated self-supervised learning for graphs. In *International Conference on Learning Representations* (2022).

[25] Hu, Z., Dong, Y., Wang, K., Chang, K.-W. & Sun, Y. Gpt-gnn: Generative pre-training of graph neural networks. In *Proceedings of the 26th ACM SIGKDD Conference on Knowledge Discovery and Data Mining* (2020).

[26] Hwang, D. *et al.* Self-supervised auxiliary learning with meta-paths for heterogeneous graphs. In Larochelle, H., Ranzato, M., Hadsell, R., Balcan, M. F. & Lin, H. (eds.) *Advances in Neural Information Processing Systems*, vol. 33, 10294–10305 (Curran Associates, Inc., 2020).

[27] Hao, B., Zhang, J., Yin, H., Li, C. & Chen, H. Pre-training graph neural networks for cold-start users and items representation. In *Proceedings of the 14th ACM International Conference on Web Search and Data Mining*, 265–273 (2021).

[28] Sun, K., Lin, Z. & Zhu, Z. Multi-stage self-supervised learning for graph convolutional networks on graphs with few labeled nodes. *Proceedings of the AAAI Conference on Artificial Intelligence* **34**, 5892–5899 (2020).

[29] Hassani, K. & Khasahmadi, A. H. Contrastive multi-view representation learning on graphs. In III, H. D. & Singh, A. (eds.) *Proceedings of the 37th International Conference on Machine Learning*, vol. 119 of *Proceedings of Machine Learning Research*, 4116–4126 (PMLR, 2020).

[30] You, Y. *et al.* Graph contrastive learning with augmentations. In Larochelle, H., Ranzato, M., Hadsell, R., Balcan, M. F. & Lin, H. (eds.) *Advances in Neural Information Processing Systems*, vol. 33, 5812–5823 (Curran Associates, Inc., 2020).

[31] Qiu, J. *et al.* Gcc: Graph contrastive coding for graph neural network pre-training. *arXiv preprint arXiv:2006.09963* (2020).

[32] Xia, X. *et al.* Self-supervised hypergraph convolutional networks for session-based recommendation. In *Proceedings of the AAAI Conference on Artificial Intelligence*, vol. 35, 4503–4511 (2021).

[33] Yu, J. *et al.* Self-supervised multi-channel hypergraph convolutional network for social recommendation. In *Proceedings of the Web Conference 2021*, 413–424 (2021).

[34] Xia, L., Huang, C. & Zhang, C. Self-supervised hypergraph transformer for recommender systems. In *Proceedings of the 28th ACM SIGKDD Conference on Knowledge Discovery and Data Mining*, 2100–2109 (2022).

[35] Du, B., Yuan, C., Barton, R., Neiman, T. & Tong, H. Hypergraph pre-training with graph neural networks. *arXiv preprint arXiv:2105.10862* (2021).

[36] Zhang, R., Zou, Y. & Ma, J. Hyper-SAGNN: a self-attention based graph neural network for hypergraphs. In *International Conference on Learning Representations* (2020).

[37] Kingma, D. P. & Ba, J. Adam: A method for stochastic optimization. *arXiv preprint arXiv:1412.6980* (2014).

[38] Zhang, R. & Ma, J. Matcha: Probing multi-way chromatin interaction with hypergraph representation learning. *Cell Systems* **10**, 397–407 (2020).

[39] Lieberman-Aiden, E. *et al.* Comprehensive mapping of long-range interactions reveals folding principles of the human genome. *Science* **326**, 289–293 (2009).

[40] Abdennur, N. & Mirny, L. A. Cooler: scalable storage for hi-c data and other genomically labeled arrays. *Bioinformatics* **36**, 311–316 (2020).

[41] Quinodoz, S. A. *et al.* Higher-order inter-chromosomal hubs shape 3d genome organization in the nucleus. *Cell* **174**, 744–757 (2018).

[42] Marchal, C. *et al.* Genome-wide analysis of replication timing by next-generation sequencing with e/l repli-seq. *Nature protocols* **13**, 819–839 (2018).

[43] Nováček, V. & Mohamed, S. K. Predicting polypharmacy side-effects using knowledge graph embeddings. *AMIA Summits on Translational Science Proceedings* **2020**, 449 (2020).

[44] Zheng, V., Cao, B., Zheng, Y., Xie, X. & Yang, Q. Collaborative filtering meets mobile recommendation: A user-centered approach. In *Proceedings of the AAAI Conference on Artificial Intelligence*, vol. 24 (2010).

[45] Harper, F. M. & Konstan, J. A. The movielens datasets: History and context. *Acm transactions on interactive intelligent systems (tiis)* **5**, 1–19 (2015).

[46] Bordes, A., Usunier, N., Garcia-Duran, A., Weston, J. & Yakhnenko, O. Translating embeddings for modeling multi-relational data. *Advances in neural information processing systems* **26** (2013).
